## Supplementary Figure for "Transcription Factor Condensates Mediate Clustering of *MET* Regulon and Enhancement in Gene Expression"

### Supplemental Figure S1

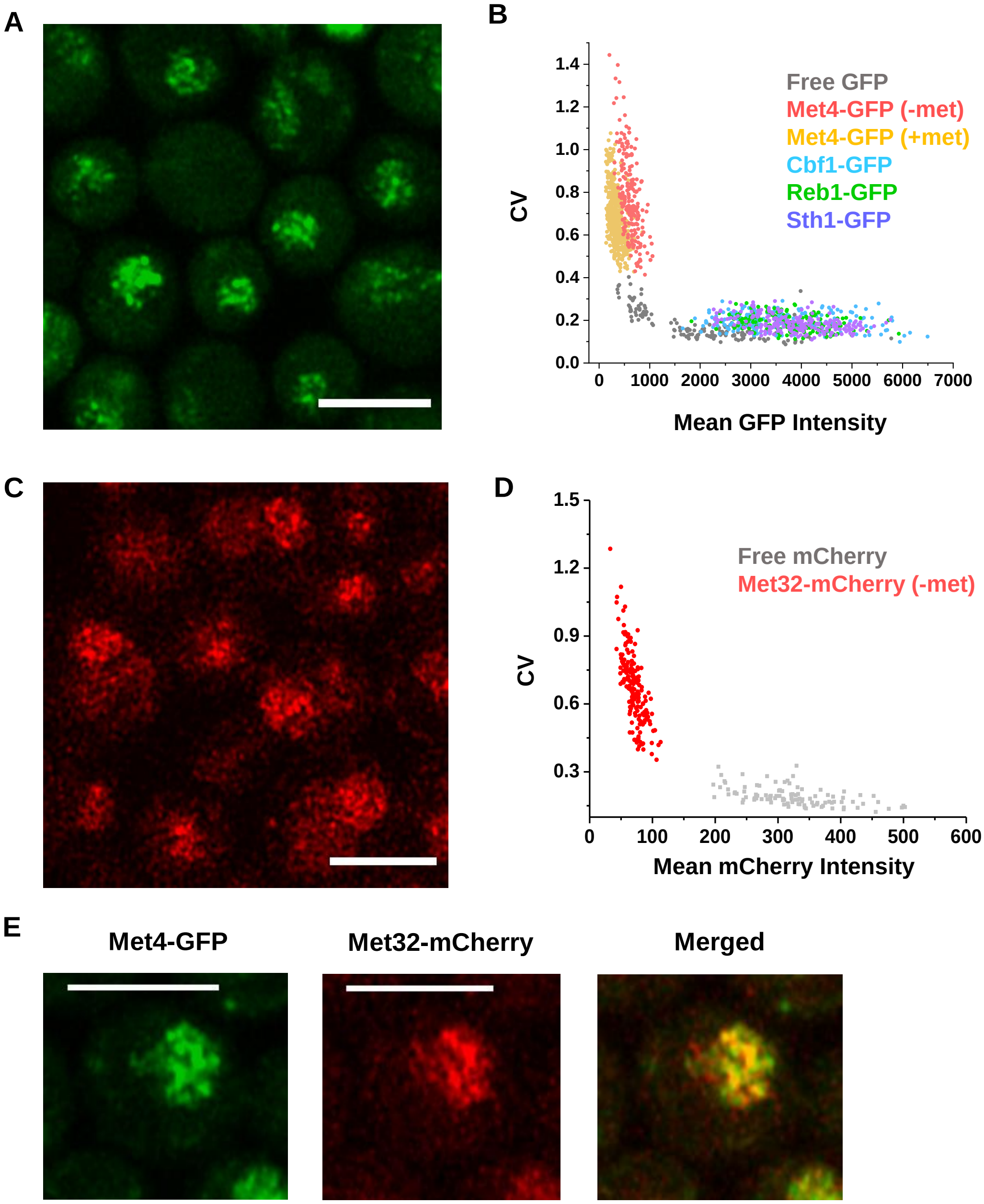


**Figure S1. A)** Additional images of Met4-GFP in live yeast cells under –met condition. Scale bars represent 4 µm. Same for below. **B)** CV of pixel intensities vs average intensity in cells expressing various GFP fusions as in Fig 1A. Each dot represents a single nucleus. CV decreases with increasing GFP concentration, and different concentrations of different factors are taken into account by the Fano number. **C)** Additional images of Met32-mCherry in live yeast cells under –met condition. **D)** Same as in B for free mCherry and Met32-mCherry. **E)** Additional images of Met4-GFP and Met32-mCherry co-expressed in live yeast cells under –met condition.

### Supplemental Figure S2

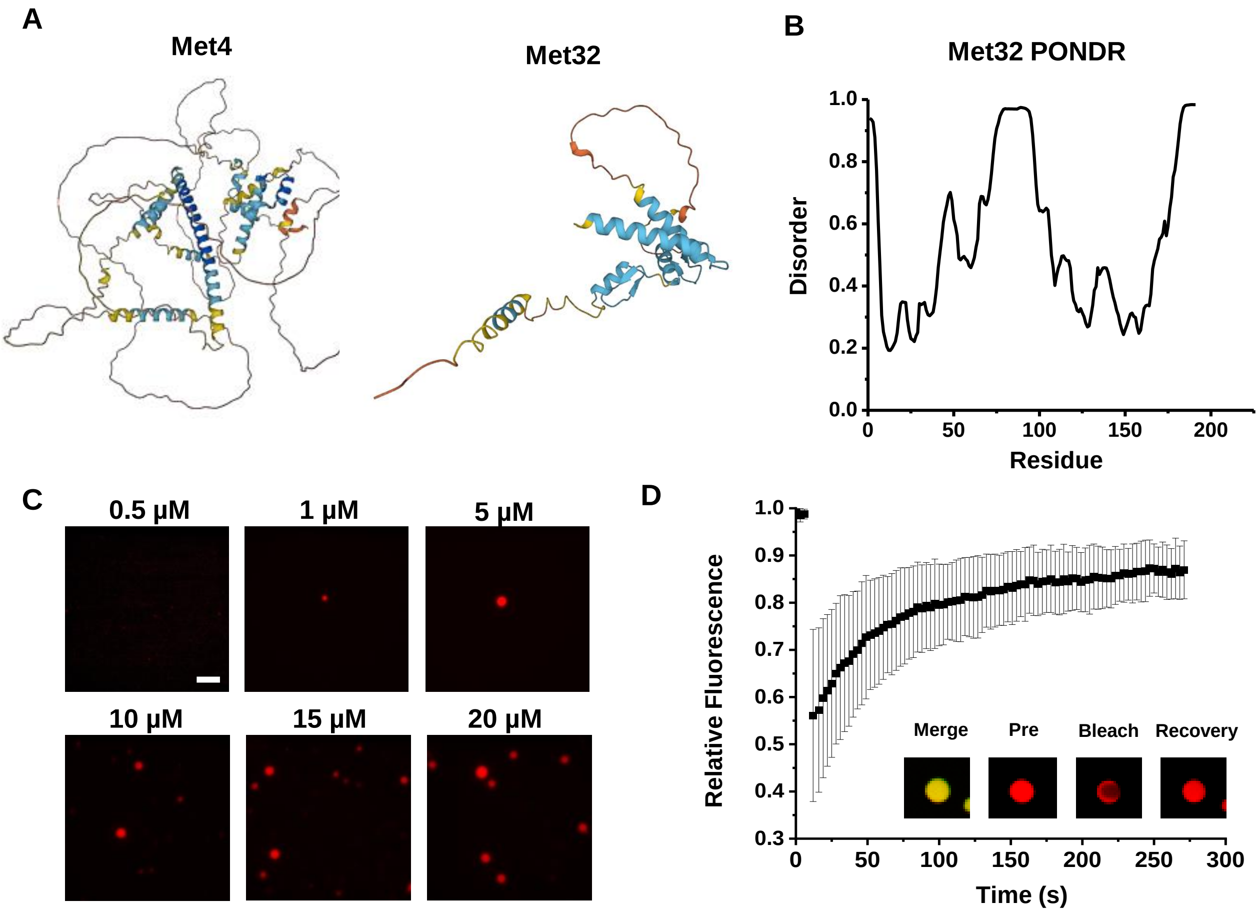


**Figure S2. A)** Predicted protein structures of Met4 (Alphafold: AF-A0A1L4AA63-F1) and Met32 (AF-A0A0L8VTL7-F1). **B)** PONDR protein disorder plot of Met32 (same analysis for Met4 is shown in Fig. 7). **C)** Met32-mCherry condensate formation at various concentrations. Purified Met32-mCherry proteins were titrated in 20mM HEPES pH 7.5, 150 mM Nacl. Scale bar represents 10 µm. **D)** FRAP curve for Met4 / Met32 co-localized droplets. 5 µM Met4-MBP and 20 µM Met32-mCherry were mixed. Intensity data collected every 3 seconds for 270 seconds.

### Supplemental Figure S3

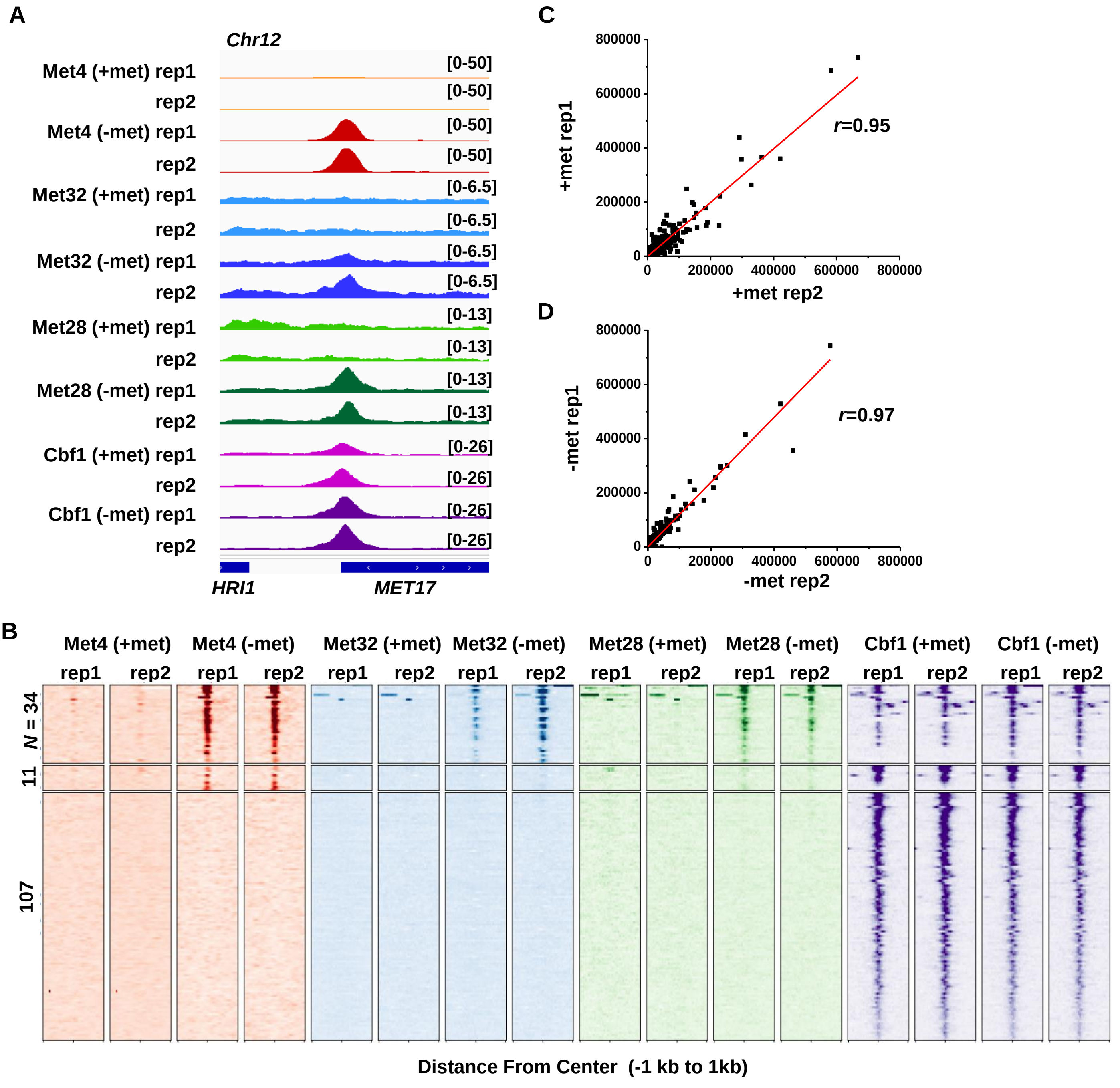


**Figure S3. A)** Additional examples of ChIP-seq data on Met TFs (Met4, Met32, Met28 and Cbf1). For each factor, two biological replicates in ±met conditions are shown. In this case, the Met TF co-binding peak is in the *MET17* promoter. **B)** Same as in Figure 3C but showing two replicates of the ChIP-seq data for the four factors in ± met conditions. **C & D)** Comparison of read counts from two biological replicates of RNA-seq data in +/- met conditions. Pearson's correlations (*r*) between the two biological replicates are shown.

### Supplemental Figure S4

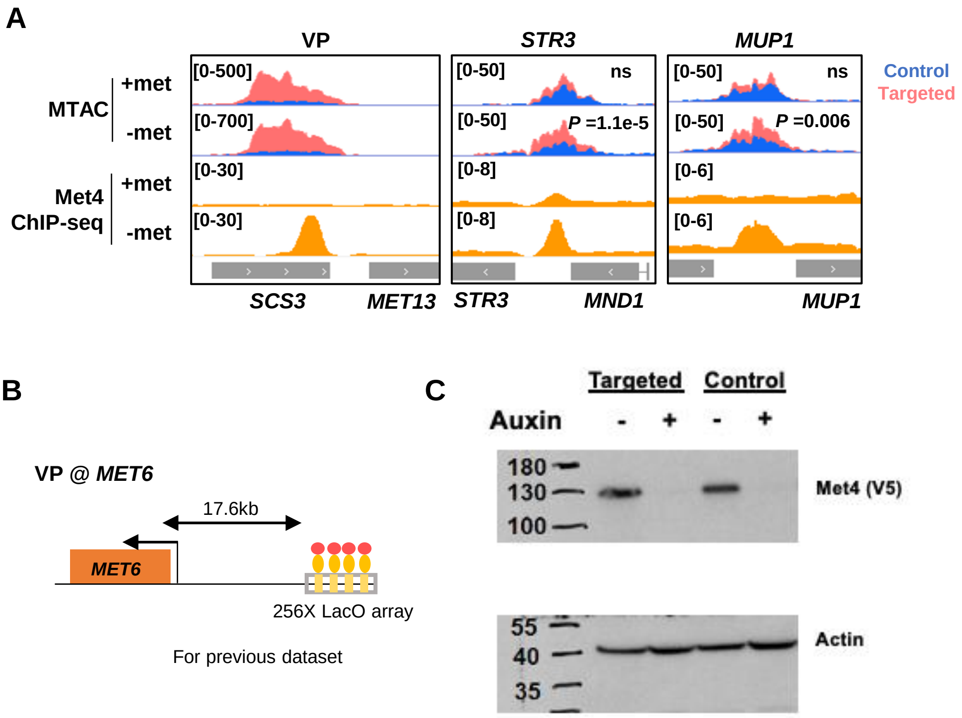


**Figure S4. A)** MTAC and Met4 ChIP-seq data at the *MET13* locus (VP), as well as at *STR3* and *MUP1* as interacting regions of the VP. MTAC signal and ChIP enrichment are both shown in ± met conditions. *P,* FDR-adjusted *P* value, Wald test by DESeq2. ns, non-significant. **B)** VP design near the *MET6* locus used for Figure 4D-F. **C)** Western blot of V5-AID-tagged Met4 ± Auxin(IAA) in MTAC targeted and control strains. Actin is probed as input loading control.

### Supplemental Figure S5

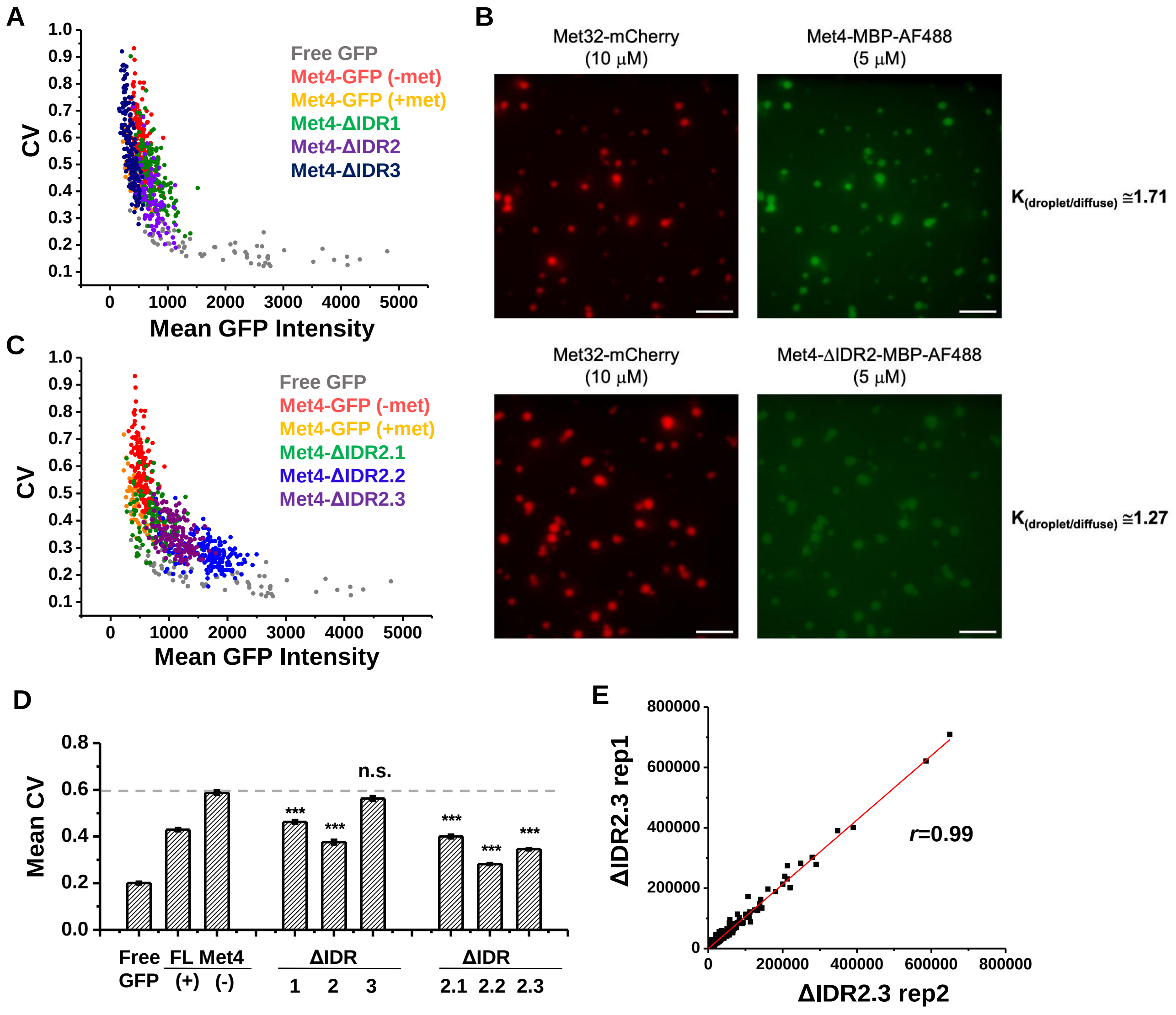


**Figure S5. A)** CV of pixel intensities vs average intensity in cells expressing free GFP or GFP fused with full length Met4 (in ± met), ΔIDR1 (aa1-69), ΔIDR2 (aa117-359), and ΔIDR3 (aa397-651). Individual dots represent single nuclei. IDR strains were grown in SCD-met. **B)** Partitioning of full length Met4 and Met4-ΔIDR2 into the Met32 condensates. The ratios of the green fluorescent intensity inside vs outside the Met32 droplet are shown on the right. **C)** Same as in A except with Met4 ΔIDR2.1 (aa117-135), ΔIDR2.2 (aa136-270), and ΔIDR2.3 (aa271-359). IDR strains were grown in SCD-met. **D)** The mean CV values of nuclear pixel intensities of different GFP fusions in panel A and B. P-values were calculated against the Free GFP. **E)** Comparison of read counts from two biological replicates of RNA-seq data of the ΔIDR2.3 strain in -met. Pearson's correlations (*r*) between the two biological replicates are shown.
